# A low-variance subspace underlies individual differences in resting state fMRI

**DOI:** 10.64898/2026.01.25.701594

**Authors:** Anastasia Borovykh, Max Weissenbacher, Stephanie Noble, Maxwell Shinn

## Abstract

Non-invasive whole-brain recordings of human brain activity, such as those from resting state fMRI (rs-fMRI), contain reliable individual differences across thousands of features, but it remains unknown whether this reliability can be isolated in a small number of dimensions. Here, we directly optimize test–retest reliability across many of these features and identify a low-dimensional linear subspace with very high reliability. These dimensions form personal fingerprints, allowing participants to be identified with high accuracy despite fingerprints explaining only a fraction of the total variance. Several dimensions are strongly associated with a single anatomical, demographic, or behavioral variable, and most dimensions can be predicted from the anatomical layout of cortical regions. Together, our findings suggest that stable individual signatures can be isolated from rsfMRI. These signatures reflect persistent anatomical and physiological differences, and provide a principled low-dimensional basis for biomarker discovery.

---

Resting state fMRI (rs-fMRI) varies both across people and within the same person over time ^1–4^. Reliable between-person differences can arise from stable traits, including lifestyle ^5,6^, intrinsic correlational structure ^7^, network organization and location ^8–10^, anatomy and brain volume ^11–16^, age ^17^, and disease ^18–20^. They can also arise from transient within-person states: people may exhibit similar patterns of thought, arousal, or autonomic state each time they are in the scanner ^21–25^. Both explanations can produce reliable between-person differences and both contribute to test–retest reliability ^26–28^. Identifying reliable differences is essential for clinical applications ^29^.

Prior work has shown that rs-fMRI can identify people across scans and can be represented in a low-dimensional form ^30–38^. However, we still do not know whether test-retest reliability itself has a low-dimensional form, and if so, whether those dimensions are associated with anatomical, physiological, or demographic variation. Ideally, we want to isolate a small number of uncorrelated dimensions that maximize reliability across repeated scans.

Here, we identify a low-dimensional subspace of rs-fMRI metrics that maximizes test-retest reliability. We find that linear combinations of simple timeseries statistics—such as the mean, standard deviation, and temporal autocorrelation—can be highly reliable, with uncorrelated dimensions having an intraclass correlation coefficient (ICC) of up to 0.98. This subspace serves as a low-dimensional fingerprint for each participant, allowing individual participants to be identified from the population with high accuracy, despite explaining only a fraction of the total variance in the original space. The components have strong associations with specific anatomical and physiological variables, and different variables are associated with different components. We release our method, reliability component analysis (RCA), as a scikit-learn– compatible library. Overall, our findings demonstrate that highly reliable dimensions can be isolated from an otherwise complex rs-fMRI signal, and that these dimensions bridge the relationship between structure and function.

## Results

### Reliability of functional brain metrics

First, we show that several simple statistics of rs-fMRI timeseries are reliable. We obtained four 15-minute scans from 999 participants from the Human Connectome Project ^39,40^ acquired across two separate days. For each scan, we defined four regional timeseries feature sets and one functional-connectivity (FC) feature set. For each of 360 cortical parcels, we computed the first three temporal moments—temporal mean, standard deviation, and skewness—as well as lag-1 temporal autocorrelation, reported previously as being highly reliable ^41^. To avoid fitting RCA directly to all 64,620 FC edges, we vectorized each FC matrix and projected it onto 360 singular vectors learned using the training scans. We retained 360 scores to match the dimensionality of the regional feature sets; they explained 84.8% of FC variance in the training set and 77.4% in the test set. To quantify reliability, we used intraclass correlation coefficient (ICC), a measure of similarity within vs between groups which takes values from −1 to 1^42^. We compute the ICC across participants, yielding one ICC per feature per region (or per singular vector for FC). We find that, consistent with previous work ^41^, all of these features have highly significant ICCs (Figure 1a, grey). We compare these to other metrics previously shown to be reliable, including the measures of spatial autocorrelation SA-*λ* and SA-∞ ^41^, as well as the “nuisance” parameters of head movement ^43^ and mean white matter and cerebrospinal fluid (CSF) signal. We find that these other metrics and nuisance parameters also have high reliability. This indicates that many basic statistics derived from fMRI timeseries are preserved across different scans from the same individual.

**Figure 1:**
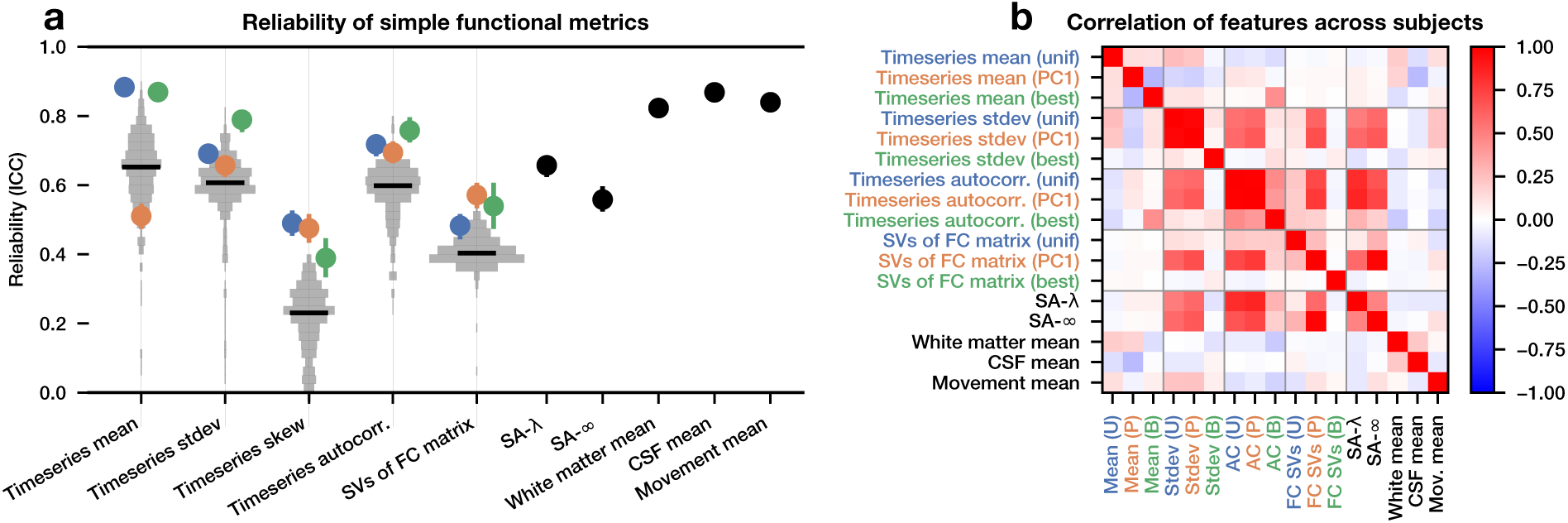
There are many reliable dimensions of rs-fMRI activity. (a) Metrics are evaluated for reliability using intraclass correlation coefficient (ICC). For metrics which produce a single value across the brain, the ICC and associated 95% confidence intervals are shown (all significant, *p* ≈ 0). For metrics which produce one value per region, a distribution of ICC for each region is shown, with a black line indicating the median. Overlaid is the ICC of the uniform projection of the regions (blue), the first principal component of the regions (orange), and the cross-validated best region (green). (b) The Spearman correlation of each of these features with each other. N=3996 scans.

If so many metrics and nuisance parameters have high reliability, does their reliability derive from the same underlying source? If so, we might expect that these metrics will be correlated across participants. We form three näıve proxies of each of the multivariate features: the average across regions (“uniform projection”), the first principal component (“PC1”), and the highest cross-validated ICC from each feature (“best region”). We compute the Spearman correlation across participants between each pair of these univariate metrics (Figure 1b) and find that for some features, the uniform projection and PC1 are correlated with each other, but for other features, they are not. Additionally, there is only a weak correlation between the “best region” projection and every other metric, despite all of them being very highly reliable on their own. This indicates that a mix of shared and independent sources of reliability can be found both within and across these features. Therefore, a specially-crafted metric combining these diverse sources of information might achieve even higher reliability than any of the näıve proxies.

### Maximizing reliability

To find a metric which maximizes reliability, we want to combine information from different brain regions into a single value for each scan. Each participant should have similar values for all of her scans, but different values from the scans of the other participants (Figure 2a). More precisely, this means finding a latent dimension in the feature space which minimizes within-participant distance but maintains a large variance across participants. To achieve this, we use a contrastive loss function, originally introduced in self-supervised machine learning ^44,45^ which found applications across neuroscience ^46–48^. Our method can perform both linear and nonlinear dimensionality reduction (see Methods), though here we focus on the linear case. We fit separate RCA models for mean, standard deviation, autocorrelation, and FC singular vector scores. We did not fit RCA to skewness because it had lower parcel-wise test–retest reliability in the initial feature survey (Figure 1a).

**Figure 2:**
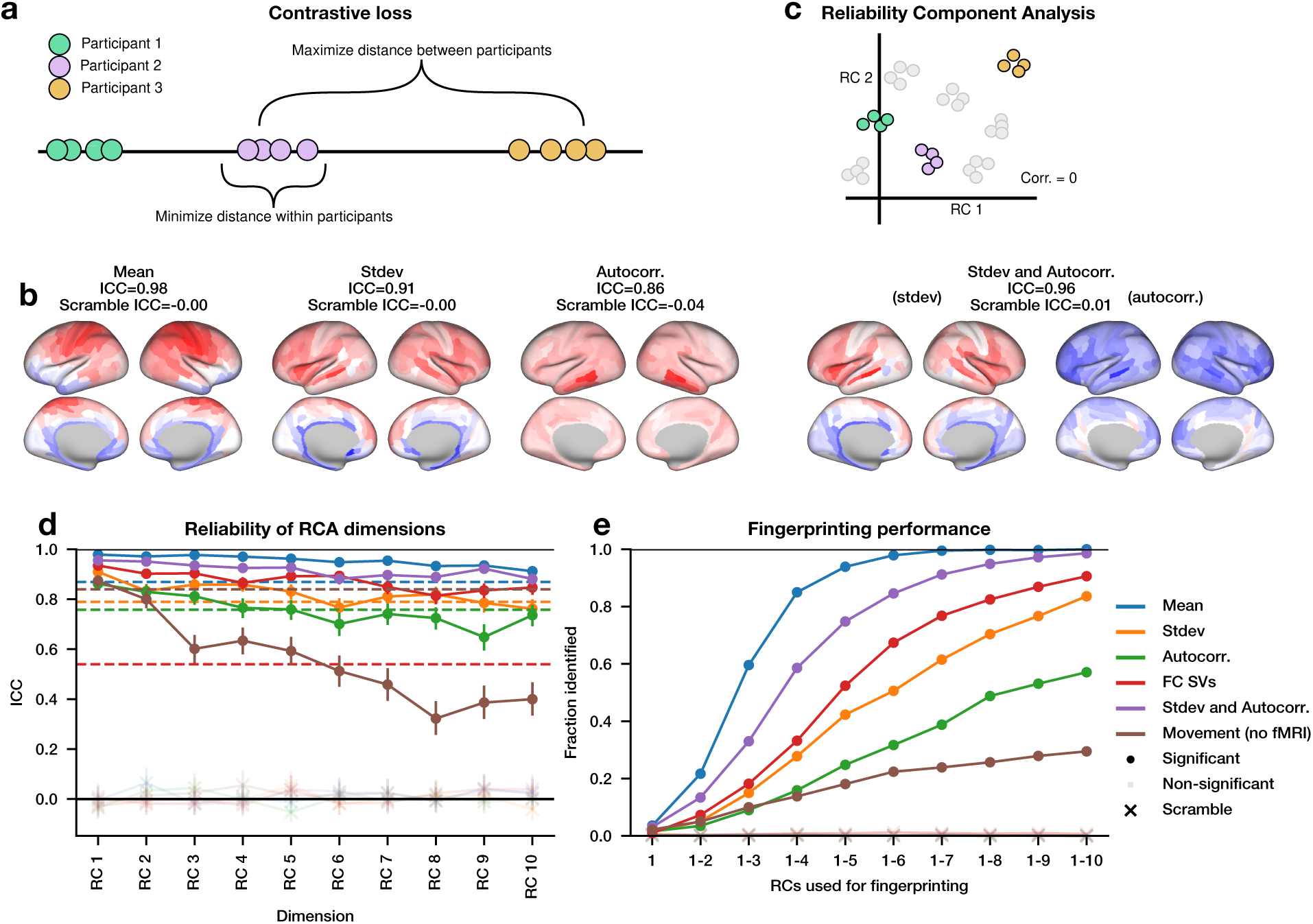
Reliability can be decomposed into linear components. (a) Schematic diagram of the contrastive loss function. (b) For each regional feature set, the surface map shows the Pearson correlation between each input feature and the first reliability component (RC) score. Intraclass correlation coefficient (ICC) for this dimension is computed using only participants in the test set. “Stdev and Autocorr.” concatenates these two feature sets and thus shows them plotted separately. (c) Schematic diagram of reliability component analysis (RCA). RCA minimizes the contrastive loss function for each dimension, sequentially under the constraint that no dimension correlates with existing dimensions. (d) The ICC for each dimension when RCA is performed on different feature sets. Points indicate ICC on the test set. For comparison, dotted lines indicate the ICC of the best cross-validated dimension from Figure 1a. “FC SVs” indicates a feature set constructed from the singular vectors of the functional connectivity matrix, and “Movement” indicates a feature set of within-scanner estimated movement parameters. Opacity indicates Bonferroni-corrected *p <* .01 across the ten RCs within each feature set; “x” indicates RCA models trained after scrambling scan–identity labels. (e) Fingerprinting performance using nearest-neighbor matching as a function of the number of included RCs. Opacity indicates Bonferroni-corrected *p <* .01 across the ten tested dimensionalities within each feature set. Models were fitted in *N* = 749 participants and evaluated in *N* = 250 held-out participants (999 total; four scans each).

Our models identified a highly reliable linear dimension for each feature set. On held-out participants, the first reliability component (RC) achieved ICCs of 0.98 for mean, 0.91 for standard deviation, 0.86 for autocorrelation, 0.93 for FC, and 0.96 for the joint standard deviation–autocorrelation representation (Figure 2b). For the regional feature sets, these values exceeded those of the best cross-validated univariate features, and the RCs had interpretable spatial patterns on the brain surface. This is in contrast to identical models fit to scrambled subject labels, which had an ICC near zero (Figure 2b). We also fit a model on a rich representation of movement, containing all estimated rotations and translations with their first temporal difference, but found only a very small increase in ICC over the total magnitude of movement alone (Movement ICC=0.840, movement dimension ICC=0.874). While we tested both linear and non-linear models, the linear models were simpler and had slightly higher reliability on the test set (Figure S1).

We next asked if combinations of different feature sets could further improve reliability. We fit a model to each pair of feature sets. We found that the model combining standard deviation and autocorrelation improved ICC substantially compared to either feature alone (Figure 2b), whereas other models produced minimal to no improvement in ICC for the most reliable feature in the pair (Figure S2). Therefore, using a more diverse set of features may, but will not necessarily, increase the amount of reliable information.

### A maximally reliable linear subspace

We showed there is a highly reliable dimension in each set of features, but are there multiple such dimensions in the same set of features? To answer this, we developed an iterative procedure for identifying uncorrelated latent dimensions that are maximally reliable, which we call reliability component analysis (RCA). In RCA, the first reliability component (RC1) is the maximally reliable latent dimension, described in the previous section. Each subsequent reliability component (RC) is the most reliable latent dimension that is uncorrelated with all other RCs. While we only enforced decorrelation within the training set, RCs tend to be uncorrelated in the test set as well (Figure S3). Taking these dimensions together, RCA forms a maximally reliable subspace, where each scan occupies a single point in the subspace, and scans from the same participant form clusters (Figure 2c).

Many reliable dimensions exist across all tested sets of rs-fMRI timeseries features (Figure 2d). These dimensions, or RCs, often correspond to meaningful spatial structure, such as interior-exterior and anterior-posterior gradients, as well as loci around the temporal lobe (Figure 2b, Figure S4).

The dimensions of each subspace form a unique functional fingerprint of the participant. We performed a fingerprinting procedure inspired by Finn et al ^30^. For each scan, we found its nearest neighbor, and then checked whether both scans were from the same participant. If so, we considered it a match. We computed the nearest neighbors in a space consisting of the first *N* RCs, with *N* ranging from 1 to 10. We found that for the Mean feature, we have near perfect fingerprinting on the test set using only 7 dimensions, and for the FC feature and the standard deviation + autocorrelation feature, approximately 90% accuracy using only the first 10 dimensions (Figure 2e). By contrast, the ICC and fingerprinting performance for the first 10 principal components are low (Figure S5). Comparable fingerprinting performance has been obtained in previous work using 35,778 FC edges ^30^ or 360^41,49^ dimensions. Note that our fingerprinting is more difficult than in Finn et al ^30^ due to a larger sample size and the lack of a target-database split; applying the correlation-based method from Finn et al ^30^ to our data gives a fingerprinting accuracy of 24.6%.

The subspaces produced by RCA are robust to differences in methodology. A non-linear variant of RCA exhibits slightly worse reliability on the test set (Figure S4, Figure S6, Figure S1), but the non-linear RCA subspaces still show substantial overlap with their corresponding linear RCA subspaces (Figure S7). We also obtained similar subspaces when fitting using different random seeds, a non-iterative variant of RCA (see Methods), and a different contrastive loss function (Figure S7).

### Reliability components are associated with biological variables

Because the RCs were reliable across repeated scans, we tested their association with stable anatomical, physiological, and demographic variables. We first tested how well the RCs predicted these participant variables. Using linear regression, we found significant held-out prediction for many anatomical, demographic, cognitive, physiological, and scan-derived participant variables (Figure 3a, filled markers). Prediction was strongest for total brain volume across the five brain-derived feature sets (test *R*^2^ = 0.36–0.58), while the movement-derived RCs strongly predicted BMI (*R*^2^ = 0.66). Due to the high predictive accuracy of brain volume, we also tested whether the predictive accuracy of these other variables might be due to their relationship with brain volume (Figure 3b, *p <* .001) ^16,50^. When we adjusted for the effect of brain volume, we still found significant but attenuated predictive power (Figure 3a, hoops).

**Figure 3:**
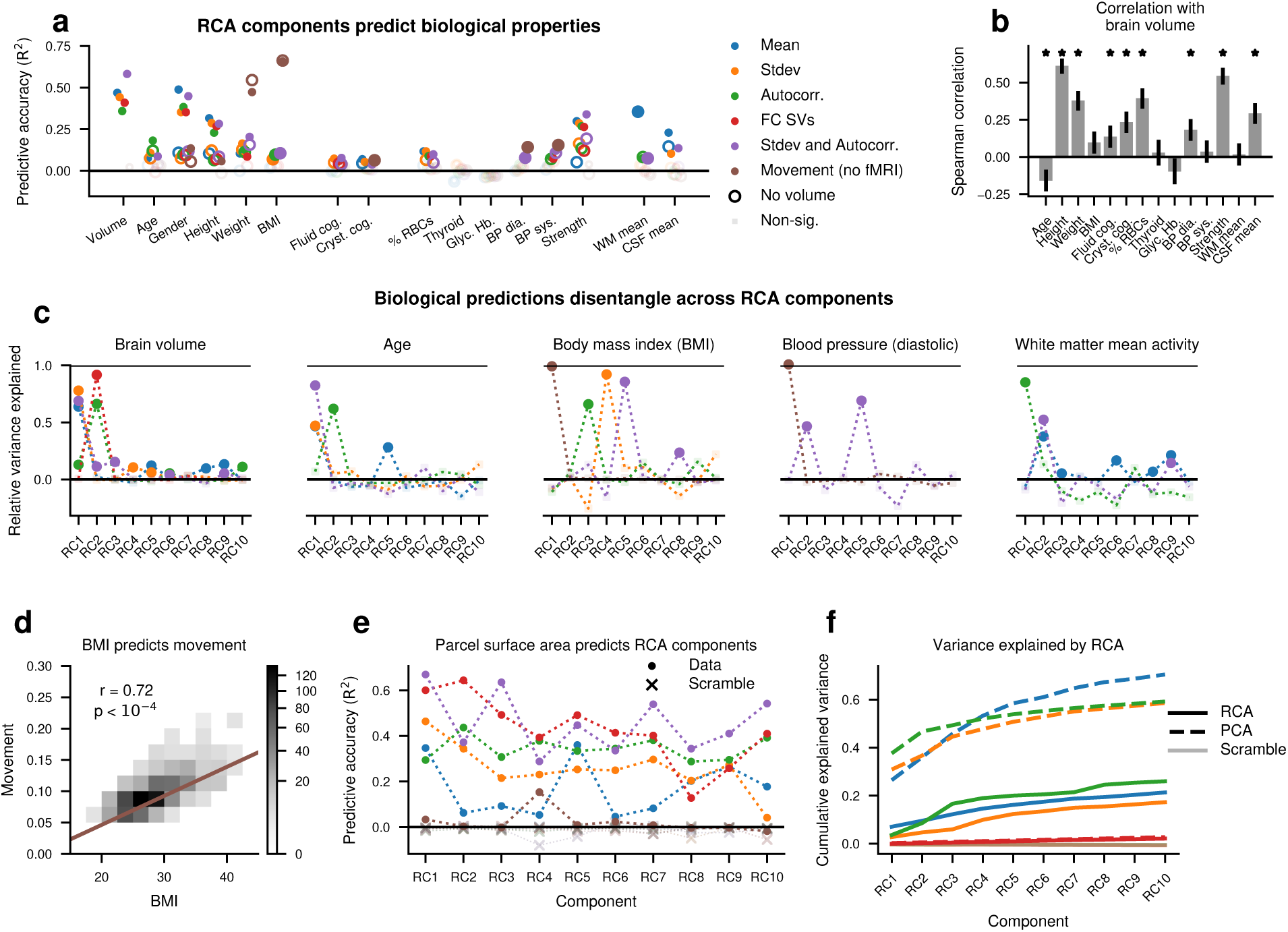
Reliability components reflect stable anatomical and physiological variables. (a) Reliability components predict anatomical, physiological, and behavioral variables on the test set (points). Y axis indicates the coefficient of determination (*R*^2^) for all variables except gender, which uses the pseudo-*R*^2^ measure of explained deviance. When brain volume is regressed out (hoops), the predictive accuracy is no longer significant for some variables. Saturated markers in (a) indicate one-sided permutation test *p <* .01 after separate Bonferroni correction across the 96 unadjusted and 90 brain-volume-adjusted predictions. (b) Spearman correlation of brain volume with each of the structural or cognitive measures from (a). In (b), * indicates *p <* .001 after Bonferroni correction across 14 correlations. (c) Fraction of the ten-RC model’s held-out *R*^2^ recovered by each one-RC model. Saturated markers indicate one-sided permutation test *p <* .01 after Bonferroni correction across ten RCs within each combination of participant variable and feature set; combinations are displayed only when the ten-RC model was significant and had *R*^2^ ≥ 0.05. (d) Correlation between body mass index (BMI) and mean movement magnitude. (e) Predictive accuracy (*R*^2^) of participant-level surface area for each parcel in predicting each RC from multiple feature sets on the test set. “x” denotes a control RCA model trained after scrambling scan–identity labels. (f) Cumulative variance explained by RCs, compared to cumulative variance explained by principal components (PCs) and by RCs trained on scrambled data, which all yield slightly negative held-out *R*^2^. Every RCA curve was significant for each dimensionality (one-sided permutation test *p <* .005).

We next asked whether this predictive power was evenly distributed across components or sparsely concentrated in a few components. We tested each individual RC’s ability to predict total brain volume, age, body mass index (BMI), blood pressure, and white matter mean signal. We found that for each model, a small number of RCs provided high predictive accuracy, while the rest of the RCs provided little to no predictive power (Figure 3c). Brain volume and age were largely predicted by the same components, and blood pressure and BMI were partially predicted by the same components, but other properties were predicted by different components. For example, the first RC for autocorrelation predicts white matter activity, the second predicts brain volume and age, and the third predicts BMI, with other autocorrelation RCs offering little to no predictive power over these variables. The high predictive power of the movement feature on BMI was largely driven by the strong relationship between BMI and total movement magnitude (Figure 3d). We reiterate that RCA had no *a priori* knowledge of anatomical, physiological, or potential confound variables, yet individual RCs strongly predicted many of these variables in held-out participants. Prediction was concentrated in one or a small number of RCs, and different variables were predicted by different RCs. This separation was not universal; for example, grip strength was predicted across several RCs (Figure S8). These results show that RCA separates reliable rs-fMRI variation into RCs that predict distinct anatomical and physiological variables.

Finally, we asked whether the reverse is true: can measures of a participant’s local brain structure predict RCs? We used participant-level measurements of parcel surface area to predict each RC, and found significant prediction for nearly all RCs (*p <* .001), especially FC and Standard deviation + Autocorrelation (Figure 3e). By contrast, the RCs formed from movement parameters and from scrambled features could not be predicted by structure (Figure 3e). Therefore, parcel size is sufficient to predict many of the most reliable dimensions of these rs-fMRI features.

One possible trivial explanation of these results is that most of the variance is reliable, and thus, RCA captures most of the variance in the signal. To test this, we compared the cumulative variance explained by RCA to that explained by PCA, an upper bound on explainable variance. We found that RCA explains only a modest amount of variance, and often the first PC explains more variance than the first 10 RCs combined (Figure 3f). This confirms that RCA specifically captures patterns of brain activity which reliably encode individual differences.

## Discussion

We showed that individual differences in rs-fMRI can be captured in a low-dimensional, low-variance linear subspace. The most reliable RCs were each strongly associated with a single anatomical, physiological, or demographic variable. In contrast to previous work ^17^, the individual variability related to age was primarily coincident with brain volume, perhaps due to the narrow range of ages among our participants. Independent of brain volume, it was most strongly represented by autocorrelation features, perhaps due to the close connection between autocorrelation and aging ^41,51^.

We performed no additional nuisance regression or global signal regression beyond the HCP minimal preprocessing pipeline and ICA-FIX denoising. Notably, we did not adjust for nuisance regressors despite their high reliability ^16,25^, because we wanted to examine the relationship of these regressors with RCs ^52^. For instance, the mean white matter signal showed a strong overlap with the reliable subspace derived from the mean on the same components as total brain volume. By contrast, the subspace derived from head movement had minimal overlap with brain-derived reliable subspaces, indicating that, even though head movement is reliable and represented in rs-fMRI activity ^43,53,54^, reliability in head movement does not drive reliability of these rs-fMRI metrics. Reliability components can include mixtures of many sources of reliable variability, including neural, anatomical, physiological, and measurement-related contributions.

Our study revealed relationships between sources of individual variability and nuisance variables. For example, the signals within the white matter and CSF were highly reliable and minimally related to brain volume, but were largely manifest in the mean, with little impact on FC. Likewise, individual variability in head movement was highly reliable but unidimensional, and this reliability was almost fully explained by BMI, a relationship known to be heritable ^55^ and causal ^56^.

An alternative interpretation of RCA is as an ideal linear subspace for clustering. Indeed, our fingerprinting performance indicates a strong clustering of participants. Unlike many linear dimensionality reduction methods, RCA does not focus on explaining variance, and indeed, explains only a modest fraction of the total variance in the data; in other words, high-variance components are not necessarily the most reliable ^57^. Nevertheless, RCA was robust across random initializations, linear and nonlinear model formulations, iterative and simultaneous component fitting, and contrastive loss functions (Figure S7), making it especially appropriate for complex datasets such as rs-fMRI, where the signal of interest is a relatively small fraction of the total variance. However, because all four scans were acquired over only two days, the reported ICCs only measure test–retest reliability over a short interval.

Not all features had many reliable dimensions. Notably, only a few RCs based on autocorrelation and movement had higher reliability than the best single feature, suggesting a simpler relationship between these features and reliability. In the case of autocorrelation, we found strong external correlates for each of the first three RCs. However, this was not the case for all sets of features. Future work with more sophisticated measures of brain structure and physiology can use RCA to further decompose reliability. For instance, nonlinear-RCA has the capacity to capture more nuanced patterns than linear-RCA, and larger datasets or data augmentation may help nonlinear-RCA capture more subtle patterns than linear-RCA. We anticipate that RCA will provide a smaller search space for functional biomarkers of slowly progressing diseases ^58,59^.

## Acknowledgments

We thank Pierre Rioux and the CBRAIN team at McGill for data access and preprocessing, and the members of the Cortical Processing Laboratory for helpful discussions. This research used the NeuroHub infrastructure and was undertaken thanks in part to funding from the Canada First Research Excellence Fund, awarded through the Healthy Brains, Healthy Lives initiative at McGill University. This research was enabled in part by support provided by Calcul Québec and the Digital Research Alliance of Canada. Data were provided by the Human Connectome Project, WU-Minn Consortium (Principal Investigators: David Van Essen and Kamil Ugurbil; 1U54MH091657) funded by the 16 NIH Institutes and Centers that support the NIH Blueprint for Neuroscience Research; and by the McDonnell Center for Systems Neuroscience at Washington University.

## Author contributions

|  | AB | MW | SN | MS |
| --- | --- | --- | --- | --- |
| Conceptualization | X |  |  | X |
| Data curation |  |  |  | X |
| Formal analysis | X | X |  | X |
| Investigation | X | X | X | X |
| Methodology | X | X |  | X |
| Project administration | X |  |  | X |
| Resources |  |  | X |  |
| Software | X | X |  | X |
| Supervision | X |  |  | X |
| Validation | X | X |  | X |
| Visualization | X | X |  | X |
| Writing – original draft | X | X |  | X |
| Writing – review & editing | X | X | X | X |

## Methods

### Data

We used data from the Human Connectome Project (HCP) ^39^. HCP data are available through the HCP repository (https://www.humanconnectome.org/study/hcp-young-adult). Users must agree to data use terms before accessing ConnectomeDB; details are provided at https://www.humanconnectome.org/study/ hcp-young-adult/data-use-terms. HCP investigators received IRB approval at the corresponding data collection sites and obtained written informed consent from all participants. The University College London and Northeastern University Human Research Protection Programs approved secondary analysis of this dataset. We accessed the data using the CBRAIN project from McGill University ^40^.

We analyzed the HCP FIX-denoised grayordinate resting state time series, registered with MSMAll and distributed with filenames of the form * Atlas MSMAll hp2000 clean.dtseries.nii. These data underwent the HCP minimal preprocessing pipeline, temporal high-pass filtering, non-aggressive ICA-FIX artifact removal, and regression of the 24 HCP motion-derived confounds ^60,61^. We parcellated the cortical data into the 360 Glasser parcels ^62^ and computed all features from the resulting time series. We performed no additional nuisance regression after the HCP processing and did not perform global signal regression. After removing incomplete scans, the dataset contained scans of 999 participants. HCP-reported gender was female for 532 participants and male for 467 participants. Participants were 22–37 years old (mean 28.7 years, SD 3.7 years).

Participants completed two resting state runs on each of two days. We treated all four runs as repeated observations, so positive pairs in the contrastive loss included both within-day and between-day run pairs. We included only participants who completed all four resting state scans, giving 999 participants. For the analyses correlating with structural and demographic variables, we excluded a small number of additional participants who refused or were unable to provide the given structural or demographic variable: N=999 for age; N=998 for height, weight, BMI, and grip strength; N=883 for fraction red blood cells (hematocrit); N=703 for thyroid-stimulating hormone; N=696 for glycated hemoglobin (glucose); and N=987 for blood pressure.

For Figure 3, we examined the full set of anatomical and physiological scalar variables in the curated HCP variable set assembled for this study: total brain volume, height, weight, body mass index (BMI), hematocrit, thyroid hormone, glycated hemoglobin, systolic and diastolic blood pressure, and grip strength. We additionally included age and HCP-reported gender as demographic variables; the NIH Toolbox fluid and crystallized cognition composites, which have high test–retest reliability ^63^; and the mean white-matter and CSF signals as scan-derived potential confounds characterized in Figure 1. Movement was analyzed as a separate feature representation, and white-matter and CSF signals were averaged across the four runs for each participant.

We split our data of 999 participants into a training set (749 participants) and a test set (250 participants) selected randomly with a fixed seed. All figures, analyses, and statistics shown are on the test set only, unless otherwise specified. The same training-test set split was used for all analyses.

Timeseries features were computed directly on the parcellated timeseries. The FC matrix singular vectors were computed by flattening the upper triangle (excluding the diagonal) of the FC matrix, and then performing SVD on the flattened matrices of the training set, using the resulting singular vectors to compute the scores for both training and test set participants. We computed the “best region” (or “best SV”) ICC for each feature by computing the ICC of each region or SV on the training set, finding the one with the maximum such value, and then reporting the ICC computed only on that region or SV within the test set.

Movement features included the estimated translation and angle (and their derivatives) in three axes, as well as the absolute magnitude, standard deviation, and temporal autocorrelation of each. They also included the mean, standard deviation, and autocorrelation of the total estimated movement magnitude timeseries, for a total of 51 values. The mean of the total estimated absolute displacement timeseries was the mean total movement magnitude.

We summarized the relationship between FC and spatial distance using the SA-*λ* and SA-∞ measures ^41^. For each scan, we binned Euclidean distances between parcel centroids in 1-mm bins, averaged FC within each bin, and fit

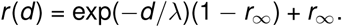

SA-*λ* is the fitted spatial decay length *λ*, and SA-∞ is the fitted long-distance asymptote *r*_∞_.

### Reliability measure

To measure the reliability across scans, we used the intraclass correlation coefficient (ICC), a measure of the variance explained by participant identity compared to the total variance. We used the strictest variant of the intraclass correlation coefficient, the one-way random-effects single-measure ICC(1,1), as our primary measure of reliability. We compute ICC following Ref ^64^ as

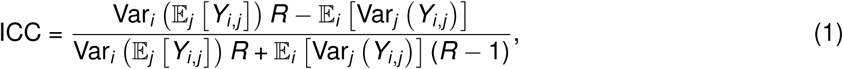

where *R* is the number of repeated observations (here *R* = 4) and *Y* is an *M* × *R* matrix, with *M* the number of participants (here *M* = 749 for the training set or *M* = 250 for the test set).

### Contrastive loss

We trained our models using a contrastive loss function ^44^. The contrastive loss function is designed to ensure that scans of the same participant (“positive” sample pairs) are located close to one another in the embedding space, while scans of different participants (“negative” sample pairs) are located far from one another in the embedding space. In other words, it optimizes for clustering positive pairs. Here, we use a one-dimensional embedding space. Let the set *P*^+^ be the set of positive (*i*, *j*) pairs and the set *P*^−^ be the set of negative (*i*, *j*) pairs, such that (*i*, *j*) ∈ *P*^+^ indicates scans *i* and *j* are from the same participant, and (*i*, *j*) ∈ *P*^−^ indicates they are from different participants. If we denote the vector of metrics from all brain regions of a particular scan by *x_i_* and the model under consideration by *f*, then our contrastive loss is given by

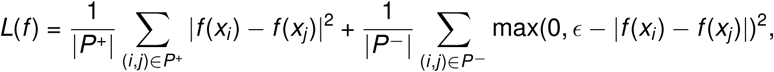

where the first sum averages pairs of scans from the *same* participant and the second sum averages pairs from *different* participants. The margin *ɛ* primarily fixes the otherwise arbitrary scale of the one-dimensional embedding: for an unregularized linear map, rescaling both the embedding and the margin leaves the optimized direction unchanged, while ICC, correlation, and nearest-neighbor ordering are invariant to a common scale factor. We therefore used *ɛ* = 1.0 as a scale convention rather than selecting it on the held-out test set.

If the training set contains *M* participants with *K* ≪ *M* scans each, we have in total *MK* (*K* − 1)*/*2 ≈ *MK* ^2^*/*2 positive pairs and *M*(*M* − 1)*K* ^2^*/*2 ≈ *M*^2^*K* ^2^*/*2 negative pairs. Therefore, there are approximately *M* = 749 more negative sample pairs for each positive sample pair, creating a large asymmetry; we thus take the expectation of the positive and negative losses separately and with an optional weighting parameter to control for the differences in magnitude.

In practice, the loss is computed over a certain batch of scans which is randomly sampled from the training set. This batch size has been found to be a key limiting factor for performance of contrastive learning methods ^65,66^; using variations such as the triplet loss ^45^ or using larger batch sizes tends to lead to better performance. The small size of the dataset used in the present study permitted the use of the entire training set in each batch, which in our experiments was essential to ensuring model training converged.

The model *f* is learned using gradient descent with the AdamW optimizer ^67^. We use a generous number of 2000 epochs with a learning rate of 10^−3^ to reach convergence on linear models, and 4000 epochs with learning rate 2 × 10^−4^ on non-linear models.

To verify robustness, we also ran simulations using an InfoNCE-style loss function, an alternative loss function frequently used in contrastive learning, given by

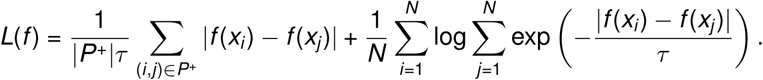

Unlike the contrastive loss above, this InfoNCE loss does not write the negative pairs as a separate *P*^−^ sum. For each scan *i*, the second term compares it with every candidate scan *j*; candidates from other participants are the negative examples.

Typically, the similarity score is computed as the cosine similarity or dot product, scaled by a temperature *τ*. In our case the output *f* can be one-dimensional, so we use the norm instead. This loss is motivated by a cross-entropy objective: for a base embedding *f* (*x*), pick the *f* (*x*^′^) such that it belongs to the same class, i.e., make the second element in the positive scan pair stand out compared to negative pair elements.

### Reliability Component Analysis (RCA)

Reliability Component Analysis is a novel technique to extract a maximally reliable linear or nonlinear subspace from a set of features, such as timeseries statistics of fMRI scans. The technique is akin to principal component analysis (PCA), but instead of iteratively extracting orthogonal dimensions of greatest variance, we extract uncorrelated dimensions of greatest test-retest reliability. The first component of an RCA subspace is the most reliable linear or nonlinear combination of the features, in the sense that the contrastive loss is minimized. For each subsequent component, we add a term to the loss function penalizing correlation with the other, already found, dimensions.

Note that RCA diverges conceptually from PCA in two important ways. First, we make components uncorrelated rather than orthogonal. This is due to the shift-invariance of the contrastive loss function— correlation is invariant to this shift, whereas the dot product is not. Second, we penalize correlation among scans (scores) in the training set, rather than in the weights (loadings). This ensures maximum separability between participants in the latent space, and also makes the technique general for both linear and non-linear models. While this makes components no longer perfectly uncorrelated in the test set, we find components are near-uncorrelated in the test set; weights are nearly uncorrelated as well, despite not optimizing for this directly in the training set or test set (Figure S3).

The input to the RCA algorithm is a dataset *D* = (*x*_1_, · · ·, *x_N_*) of feature vectors *x_i_* ∈ R*^F^* from each scan of each participant, where features here are the regional mean, standard deviation, or autocorrelation of the timeseries; the singular vectors of FC; movement parameters; or combinations thereof. RCA also requires a symmetric binary matrix *s* of size *N* × *N*, where an element *s_i_*,*_j_* = 1 if scans *i* and *j* come from the same participant, and 0 if they do not. In the language of contrastive learning, this groups pairs of feature vectors (scans) into positive (*s* = 1) and negative (*s* = 0) pairs depending on whether they belong to the same participant.

To fit RCA components, we fit a function *f*: R*^F^* → R for each component which optimizes the contrastive loss function. We write *f* (*D*) = (*f* (*x*_1_), *…*, *f* (*x_N_*))^T^ ∈ R*^N^* for the vector of scores obtained by applying *f* to all scans. In the case of the standard linear version of RCA, *f* is a linear model, i.e., *f* (*x_i_* |*w*) = *x*^T^*w*, so *f* (*D*|*w*) = *Dw*, for a weight vector *w*. For the nonlinear version, *f* is a small neural network, consisting of 10 hidden ReLU units.

For the first component *k* = 1, we initiate the computation of RCA components by fitting the model to the dataset *D*,

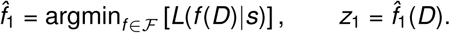

For higher components *k >* 1, we add a penalty term to assert decorrelation with earlier components. We note that our loss function is invariant to differences in absolute magnitudes of the embeddings, i.e., a constant offset would not change the loss function. Therefore, we enforce decorrelation, which is also invariant to these differences in mean, rather than orthogonality, which is not. We assert this decorrelation on the training set only, namely,

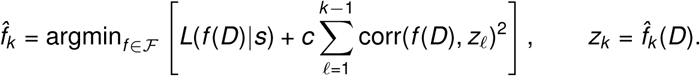

Here, *z_k_* is the vector of scores for component *k* on the training dataset, and *c* is a constant. For the results shown here, we used *c* = 0.1.

In practice, the intractable arg min is replaced with a certain number of gradient descent steps to minimize the loss, chosen so that convergence is achieved for the contrastive loss. By repeating the procedure outlined above *d* times, we obtain *d* RCA dimensions.

We also tested a non-iterative variant of RCA which fit all components simultaneously. Without imposing orthogonality or decorrelation explicitly, we rely on an observation from deep learning ^68^: different factors become disentangled in the representations learned by the model due to some implicit regularization enforcing each direction to extract different features from the data. We fit at most four components instead of ten (as we observed a rapid drop in the reliability of individual components as we increased beyond four components). Model fitting was accomplished in the linear case by making *w* a *F* × 4 matrix instead of a *F* × 1 vector, and in the nonlinear case by letting *f* (*D*) return a *N* × 4 matrix instead of a *N* × 1 vector. The rapid drop in reliability is likely related to ceiling effects, as components are already near ICC of 1.0.

RCA differs conceptually from canonical correlation analysis (CCA) and partial least squares (PLS) by using the same projection matrix on both sides and by supporting more than two observations. It differs conceptually from linear discriminant analysis (LDA) by allowing many classes to occupy the same linear dimension, enforcing clustering rather than linear separability. It shares similar goals to some eigenvector-based approaches for maximizing reliability ^69–71^ but includes the critical nonlinear term in the loss function, which precludes a closed-form solution using SVD or other eigenvector-based methods.

### Visualization

To visualize an RC, we computed the Pearson correlation across held-out test scans between its score and each parcel-level input feature. For example, for an RC derived from lag-1 autocorrelation, each map value is the correlation between that parcel’s lag-1 autocorrelation and the RC score. We display feature–RC correlations rather than linear weights because the regional inputs are strongly correlated and the weights are unregularized. We also chose this procedure so that the same visualization could be used for nonlinear RCA, which does not have a single linear weight vector.

### Structural and demographic prediction

For each participant, we averaged each RC score across the four scans. We used unregularized linear regression to predict each continuous participant variable from the first ten RC scores, fitting on the training participants and evaluating on the held-out test participants. For each test, we generated 12,000 null values by permuting the correspondence between the participant-level RC score vectors and the participant variable and repeating the same training/test procedure. To remove the linear association with total brain volume, we also regressed brain volume from the variable of interest and predicted the residual as above.

Results were evaluated with the coefficient of determination (*R*^2^), namely,

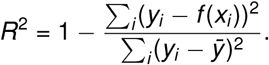

Gender was predicted with logistic regression. To regress out brain volume, we performed linear regression of brain volume onto the independent variable first and then performed logistic regression on the residual. Results were evaluated using explained deviance definition of pseudo-*R*^2^, namely,

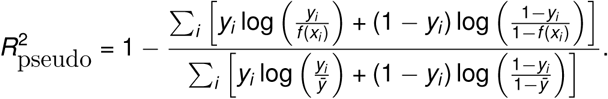

For each RC, we fit a separate one-predictor model in the training participants and evaluated its *R*^2^ in the held-out test participants. We divided this value by the held-out *R*^2^ of the corresponding ten-RC model to measure the fraction of full-model prediction recovered by that RC. We generated 1,200 participant-level permutations for each one-RC test. To avoid unstable ratios, we displayed only combinations of participant variable and feature set for which the ten-RC model was significant and explained at least 5% of held-out variance.

One-sided empirical permutation p values were computed as 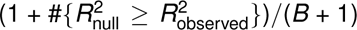, where *B* is the number of permutations. In Figure 3a, Bonferroni correction was applied separately across the 96 unadjusted predictions (16 participant variables by six feature sets) and the 90 brain-volume-adjusted predictions (15 participant variables by six feature sets). In Figure 3c and Figure S8, correction was across the ten RCs within each combination of participant variable and feature set. In Figure 3b, correction was across the 14 reported correlations with brain volume.

To predict RCs from brain-wide measures of structure, we used ridge regression to predict the first 10 RCs from the surface area of each of our 360 parcels. We performed 5-fold cross validation within the training set to select the best penalty constant, and then fit the best of these again on the training set, evaluating fit on the test set.

To comply with the HCP reporting guidelines in displaying the relationship between BMI and movement in Figure 3, we set all histogram bins with fewer than three participants to white for the visualization only.

Since RCA components are not orthogonal, we estimated explained variance using regularized linear regression, fitting on the training set by predicting the features using the components, and then evaluating variance explained on the test set. To ensure a fair comparison, the same procedure was used for estimating variance with the PCA components.

These permutations of participant variables differ from the scrambled-label controls in Figures 2 and 3e, for which RCA itself was trained after shuffling the association between scans and participant identity.

## Data and code availability

All data are from the Human Connectome Project young adult 1200 subject release ^39^. Access to these data is controlled by HCP; qualified researchers must agree to the applicable data use terms before obtaining access through ConnectomeDB. Code to perform RCA is available as the scikit-learn–compatible Python package scikit-rca, installable from pip, or from the URL https://github.com/maxweissenbacher/scikitrca. Code to reproduce the figures is available at https://github.com/mwshinn/figures_for_rca_paper.

## Supplemental figures

**Figure S1:**
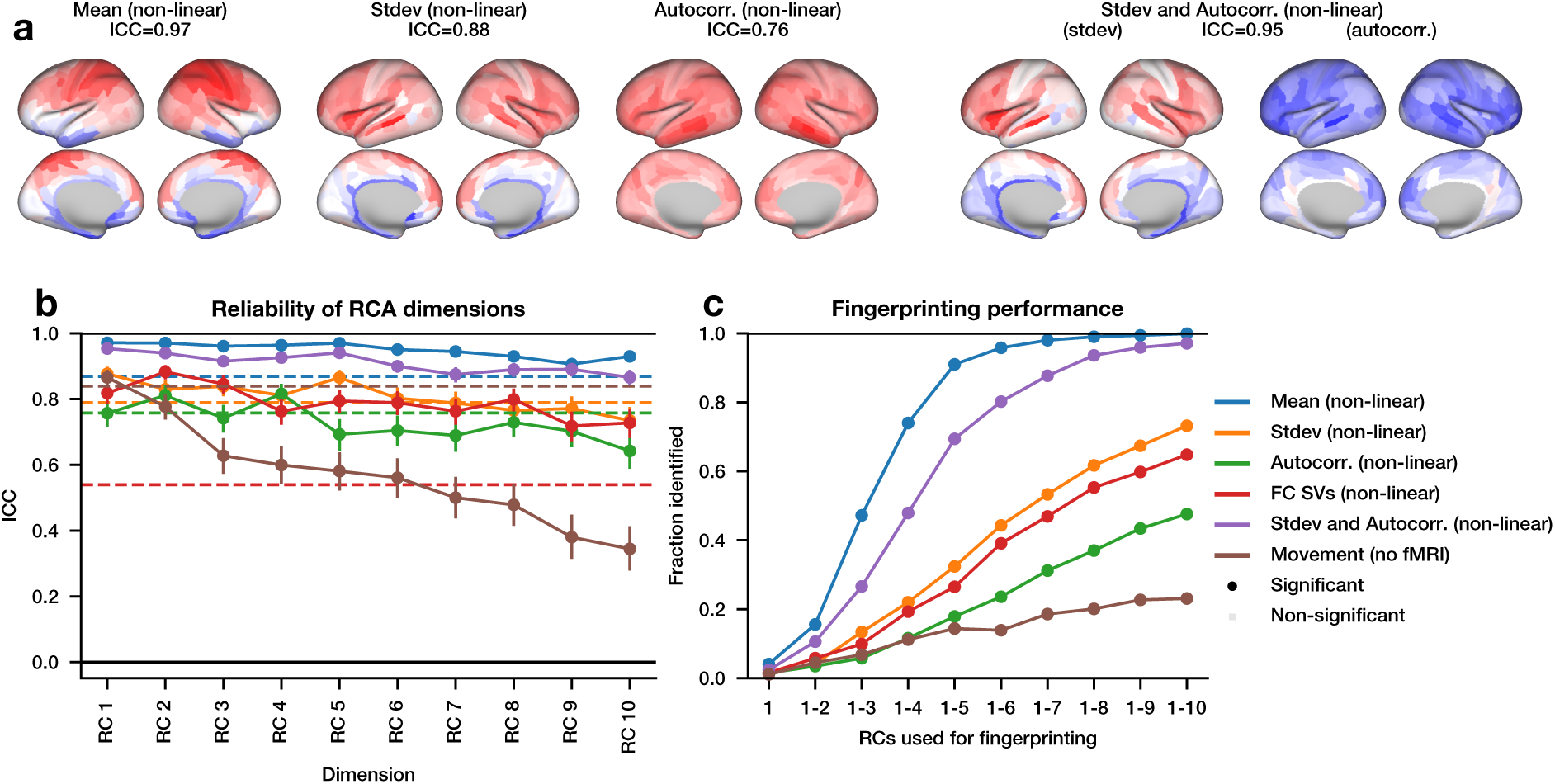
Nonlinear RCA is slightly outperformed by linear RCA. The analyses match Figure 2, except that RCA used a neural network with one hidden layer of 10 ReLU units. Models were fitted in 749 participants and ICC and fingerprinting were evaluated in 250 held-out participants.

**Figure S2:**
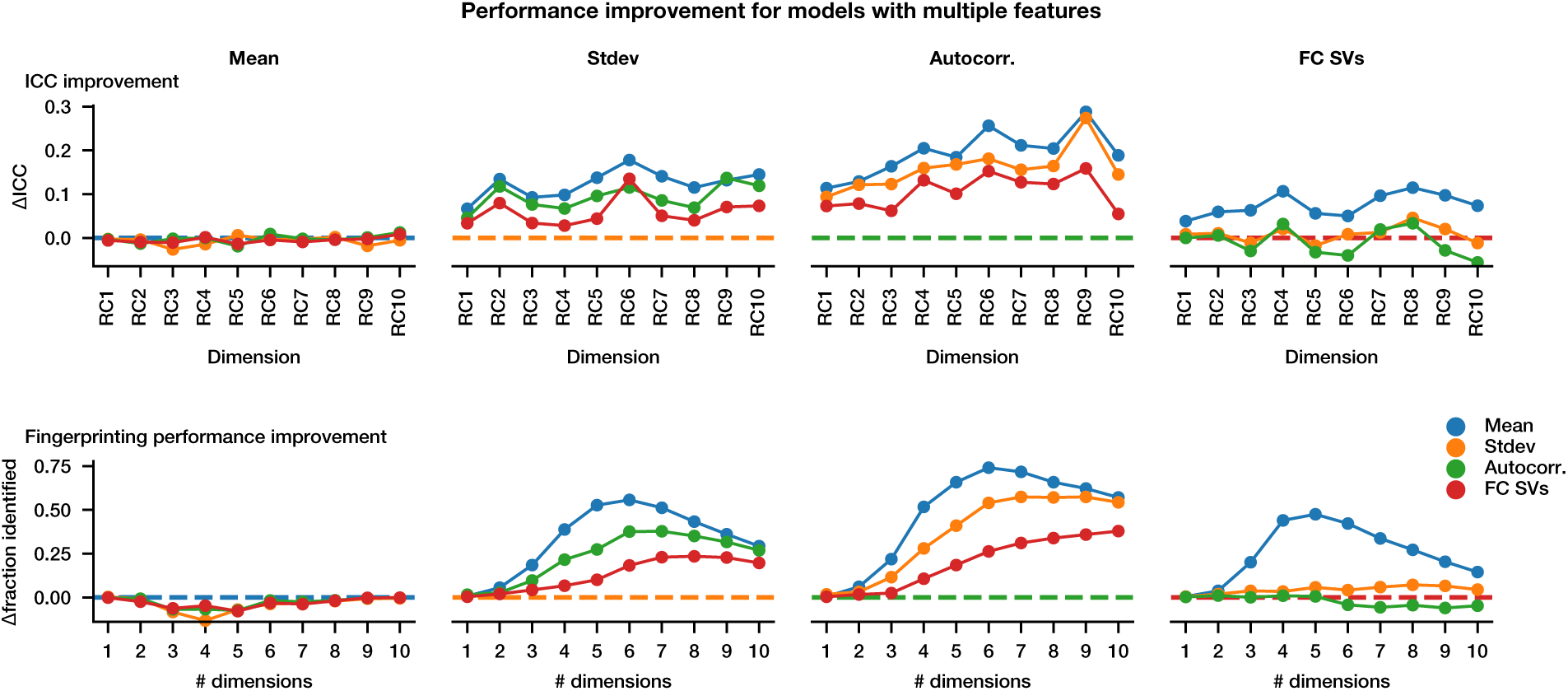
RCA performed on pairs of features. For each model, we show the difference in ICC (top) and fingerprinting performance (bottom) between a model that includes a second set of features. Positive values indicate the model is better with two sets of features instead of one. ICC and fingerprinting performance are calculated only using the test set, which was not used in training the model.

**Figure S3:**
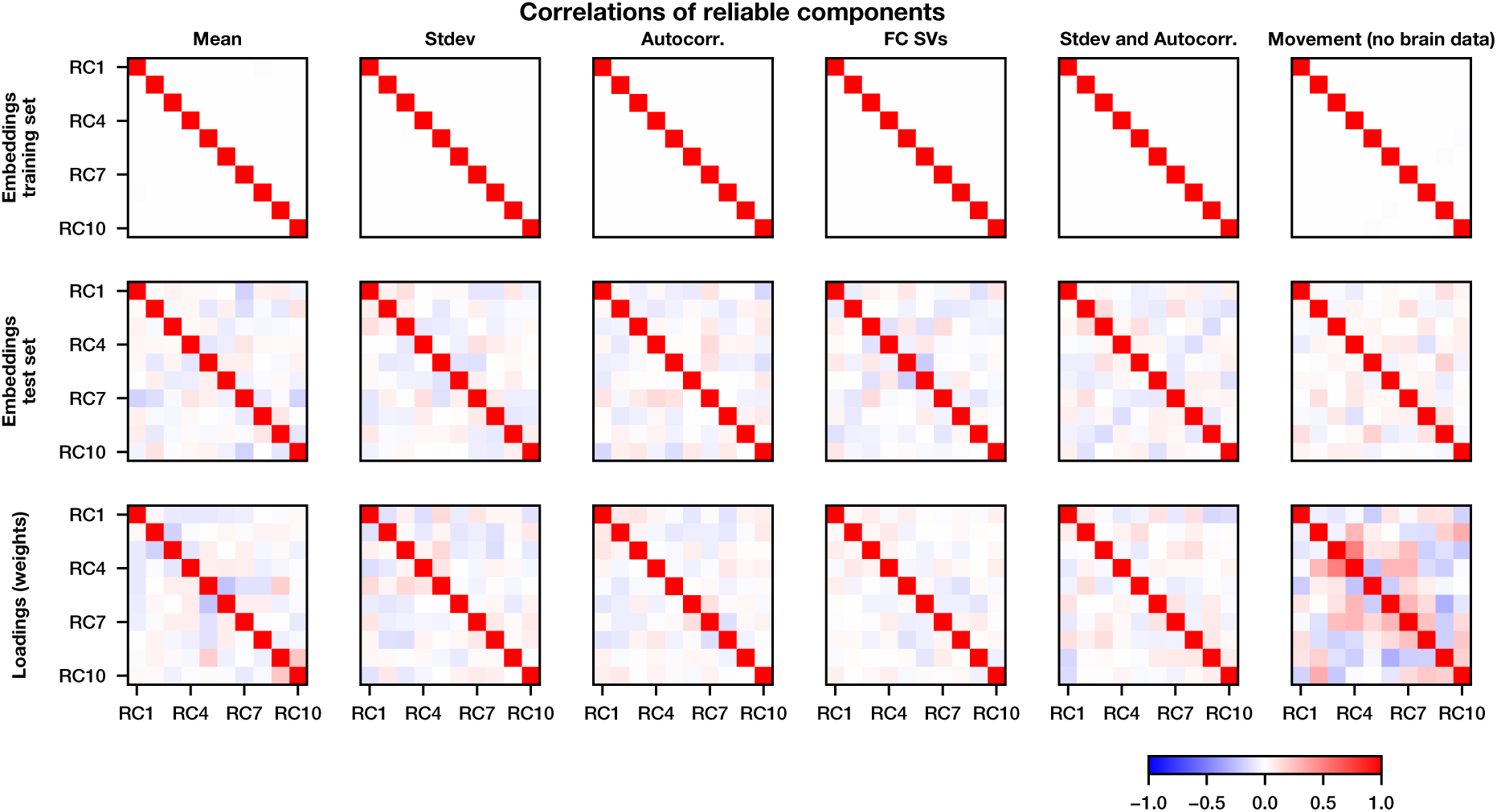
RCA components are nearly uncorrelated on the test set. RCA constrains components to be uncorrelated in the training set, but does not constrain the test set or weights to be uncorrelated. The first two rows show Pearson correlations between RC scores across all training scans and all test scans, respectively; the third row shows correlations between the linear loading vectors.

**Figure S4:**
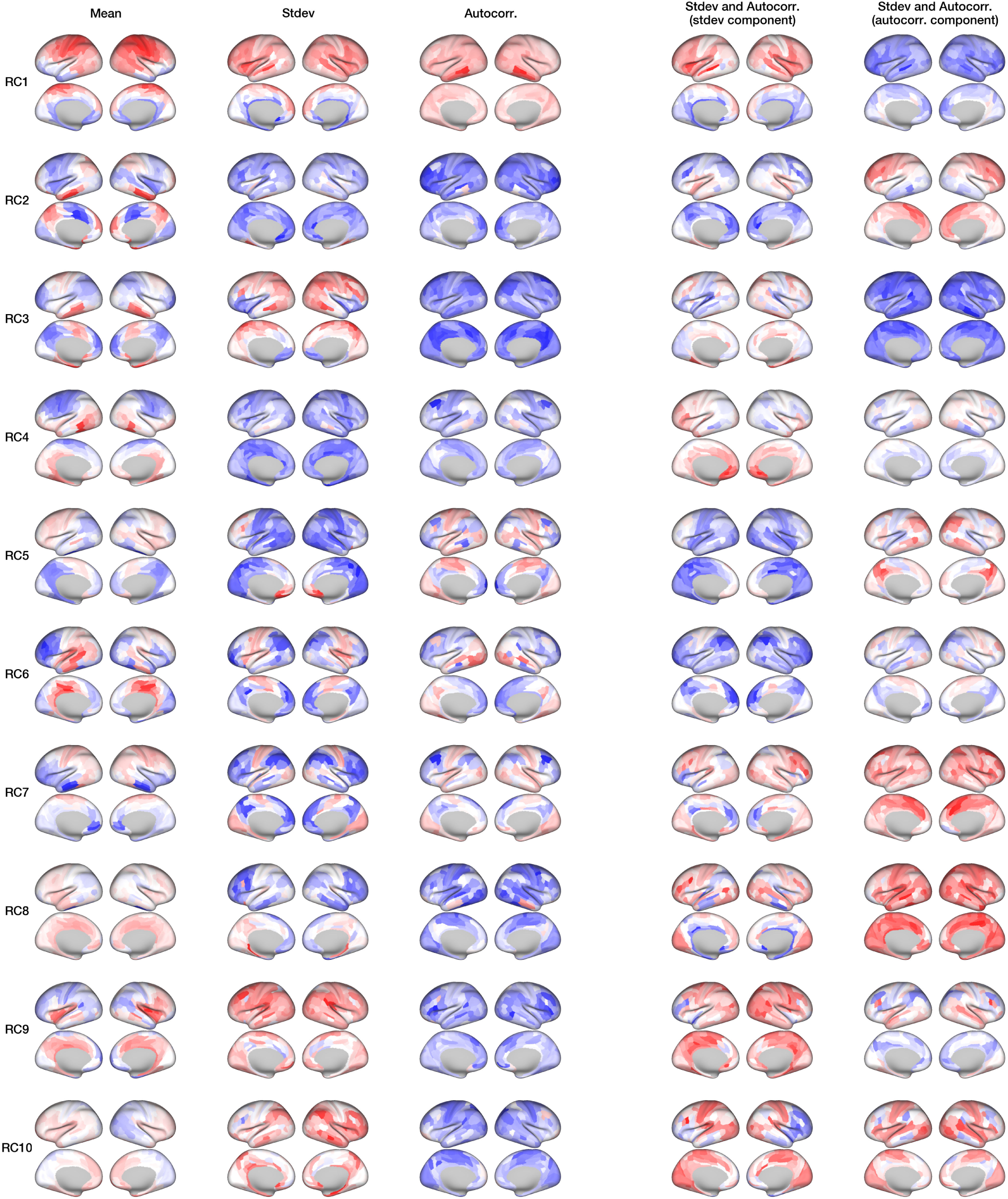
The first 10 components in RCA. For each of the four metrics, we show the first 10 components of RCA. For joint standard deviation and autocorrelation, we show the RCA components of each of these separately. Each surface is the held-out feature–RC correlation map defined in the Visualization Methods.

**Figure S5:**
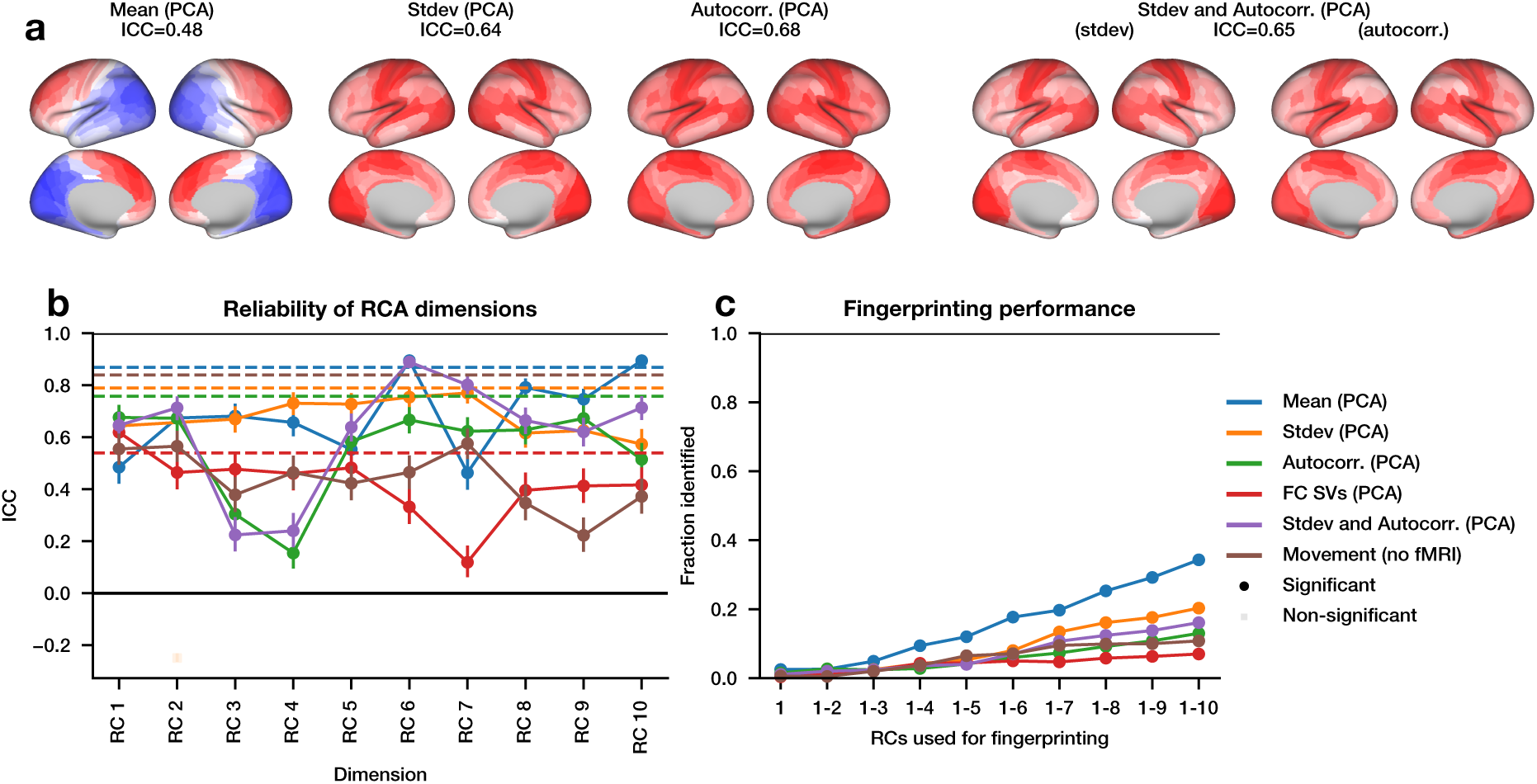
PCA does not produce reliable dimensions. PCA was fitted using the training scans and then applied to both training and held-out test scans before the same ICC and fingerprinting evaluation used in Figure 2.

**Figure S6:**
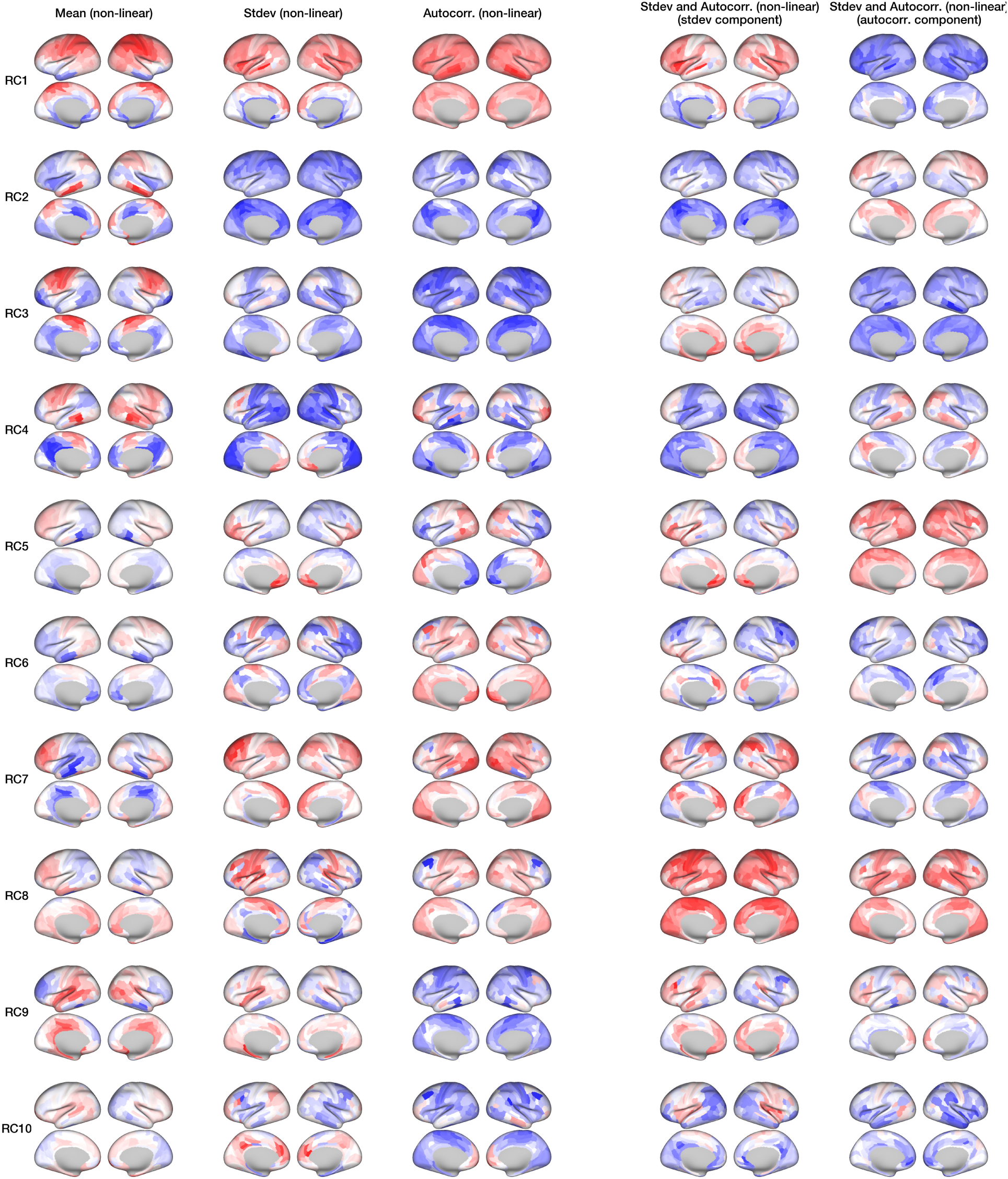
The first 10 components in nonlinear-RCA. For each of the four metrics, we show the first 10 components of nonlinear-RCA. For joint standard deviation and autocorrelation, we show the nonlinear-RCA components of each of these separately. As in Figure S4, each surface shows a held-out feature–RC correlation.

**Figure S7:**
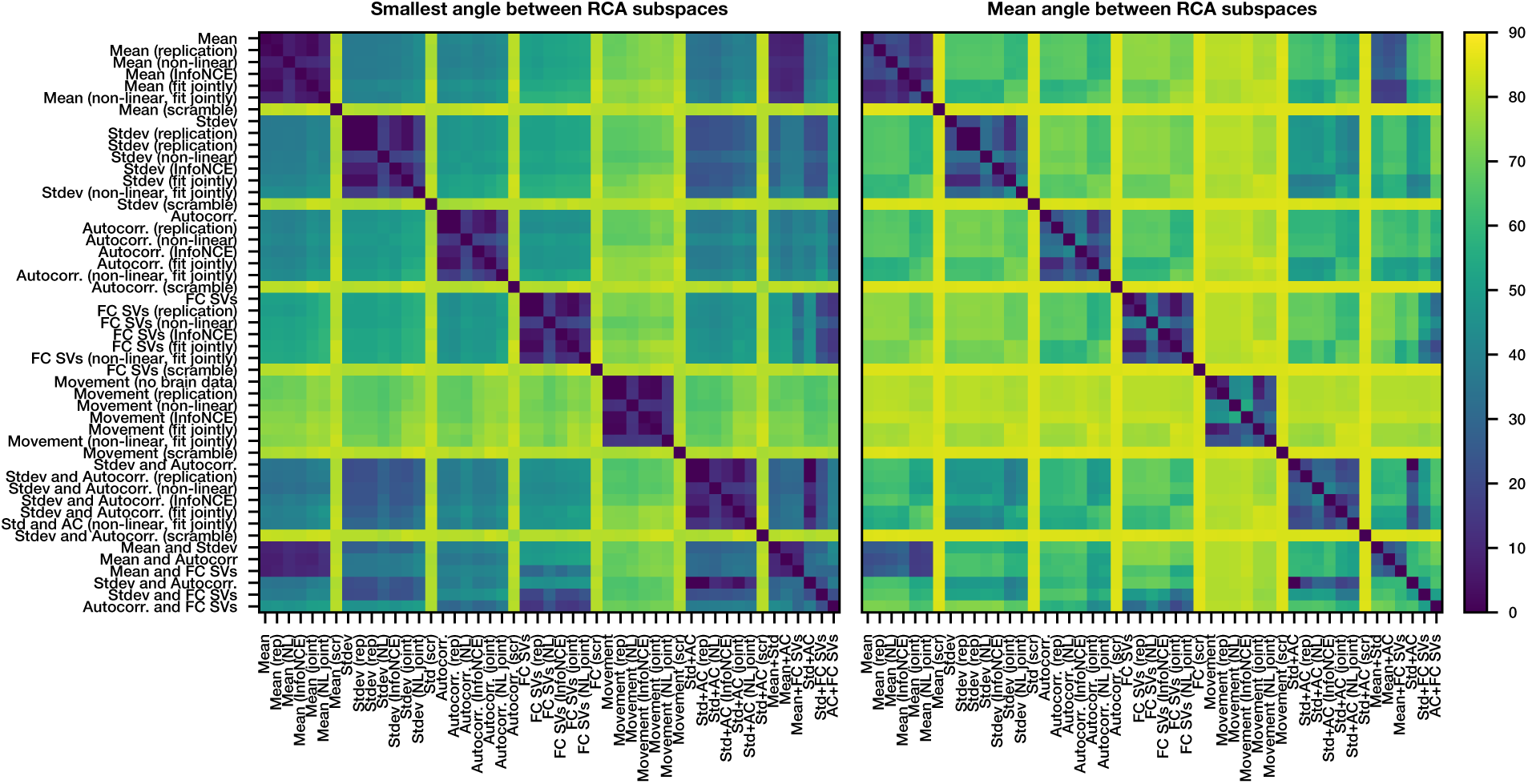
Differing overlap between reliable subspaces. Subspaces were fitted to each feature set using the primary model, a repeat fit with a different initialization, iterative nonlinear RCA, jointly fitted linear and nonlinear variants, and the InfoNCE loss; scrambled-label fits serve as controls. Overlap on held-out scans was summarized by the smallest principal angle (left) and the mean principal angle (right) across the shared dimensions of each pair of subspaces.

**Figure S8:**
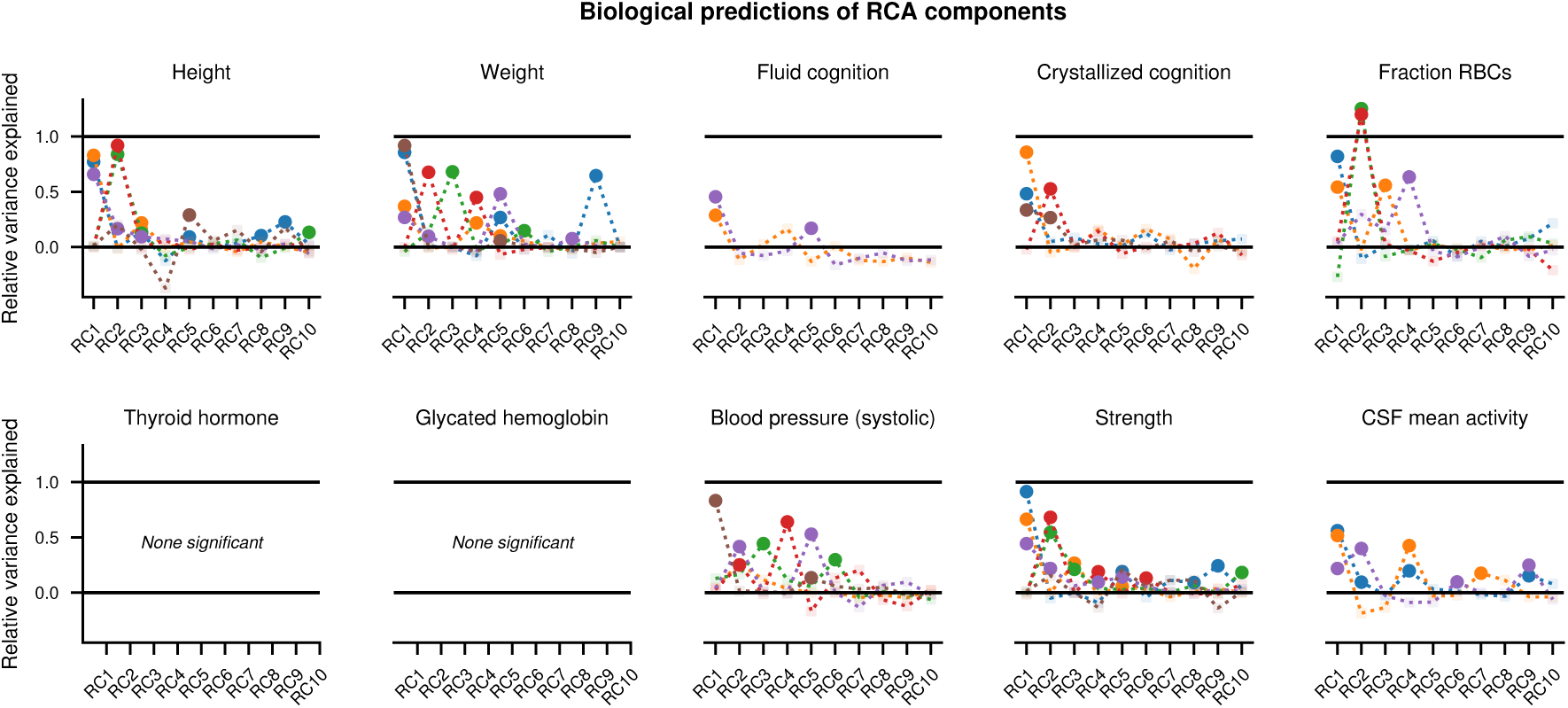
Predictions of biological variables from RCs. Same calculation as Figure 3c for the remaining reported participant variables. Each one-RC regression was fitted in the training participants and evaluated in held-out participants; opacity indicates one-sided permutation *p <* .01 after Bonferroni correction across the ten RCs within each combination of participant variable and feature set.

